# Visual uncertainty unveils the distinct role of haptic cues in multisensory grasping

**DOI:** 10.1101/2022.01.25.477675

**Authors:** Ivan Camponogara, Robert Volcic

**Author notes:** **Contributions:** I.C. and R.V. designed research. I.C. performed research. I.C. and R.V. analyzed data. I.C. and R.V. wrote the paper.

## Abstract

Multisensory grasping movements (i.e., seeing and feeling a handheld object while grasping it with the contralateral hand) are superior to movements guided by each separate modality. This multisensory advantage might be driven by the integration of vision with either the haptic position only or with both position and size cues. To contrast these two hypotheses, we manipulated visual uncertainty (central vs. peripheral vision) and the availability of haptic cues during multisensory grasping. We showed a multisensory benefit irrespective of the degree of visual uncertainty suggesting that the integration process involved in multisensory grasping can be flexibly modulated by the contribution of each modality. Increasing visual uncertainty revealed the role of the distinct haptic cues. The haptic position cue was sufficient to promote multisensory benefits evidenced by faster actions with smaller grip apertures, whereas the haptic size was fundamental in fine-tuning the grip aperture scaling. These results support the hypothesis that, in multisensory grasping, vision is integrated with all haptic cues, with the haptic position cue playing the key part. Our findings highlight the important role of non-visual sensory inputs in sensorimotor control and hint at the potential contributions of the haptic modality in developing and maintaining visuomotor functions.

## Introduction

A large proportion of grasping actions are directed toward objects we can sense with multiple modalities. For instance, when grasping with one hand an object we already hold in the other hand, the properties of the object, such as its size and position in space, are provided by both vision and haptics (touch and proprioception). The integration of these redundant sensory cues fosters a consistently superior grasping performance compared to when movements are guided by each modality alone (Camponogara & Volcic, 2019a, 2019b). Even more intriguingly, the same superior grasping performance is achieved when the haptic size cue is not provided and vision is complemented by only the haptic position cue (Camponogara & Volcic, 2021b).

The elusive effect of the haptic size cue in the multisensory integration process might result from two different causes. The superior performance in multisensory grasping might arise from the visual and haptic integration at the level of the position cues only which would reduce the uncertainty about the position of the object in space (Battaglia et al., 2010; Carey & Allan, 1996; Chen, Sperandio, & Goodale, 2018; Sperandio, Kaderali, Chouinard, Frey, & Goodale, 2013). As a consequence, the object size estimation would be solely determined by vision (Camponogara & Volcic, 2021b). Alternatively, the visuo-haptic integration might occur both at the level of the position cues and at the level of size cues, but the dominance of the more reliable visual size cue would completely overshadow the haptic size cue, making it hard to determine whether the multisensory size information is truly integrated.

The main aim of this study was to contrast these two alternative explanations by disrupting visual information during multisensory grasping. The quality of visual information was manipulated by modulating the participants’ gaze direction and, by this, the grasping actions were executed in either central (foveal) or peripheral vision. Because visual acuity sharply declines with retinal eccentricity (Rosenholtz, 2016; Strasburger, Rentschler, & Jüttner, 2011), object’s size and position estimates are noticeably impaired in peripheral compared to central vision (Baldwin, Burleigh, Pepperell, & Ruta, 2016; Bock, 1993; Brown, Halpert, & Goodale, 2005; Collier, 1931; Goodale & Murphy, 1997; Newsome, 1972; Schneider, Ehrlich, Stein, Flaum, & Mangel, 1978; Thompson & Fowler, 1980). Moreover, multisensory integration studies in perception have shown that as the quality of visual information gradually declines, the object size estimation shifts toward more haptically-based perceptual judgments (Derrick & Dewar, 1970; Ernst & Banks, 2002; Gepshtein & Banks, 2003; Helbig & Ernst, 2007; Heller, 1983; Van Doorn, Richardson, Wuillemin, & Symmons, 2010). It might be thus expected that increasing visual uncertainty through peripheral vision should let the haptic size cue effect emerge also in conditions of multisensory grasping.

With respect to movements in central vision, grasping movements in peripheral vision are generally slower, with larger grip apertures and with a poorer grip aperture scaling (Brown et al., 2005; Goodale & Murphy, 1997; Hesse, Ball, & Schenk, 2012; Schlicht & Schrater, 2007; Sivak & MacKenzie, 1990, 1992; Watt, Bradshaw, & Rushton, 2000). Introducing additional haptic cues might thus refine grasping movements in several ways depending on the contribution of haptic position and size cues. The integration of the haptic position cue would reduce the overall positional uncertainty, which would translate into faster movements and narrower grip apertures. Analogously, the contribution of the haptic size cue would diminish the uncertainty relative to the object size and would be revealed by an improved grip aperture scaling. However, if the haptic size cue is not part of the integration process, the sensitivity to changes in object size should remain unaffected.

We tested these predictions in two experiments. In the first experiment, we contrasted grasping performance under peripheral vision conditions, with (pVH) or without (pV) additional haptic cues, along with the central vision conditions (V, VH) and a haptic only (H) condition. In the second experiment, we further teased apart the contribution of haptic cues when grasping handheld objects in peripheral vision by selectively withdrawing the haptic size cue and providing the haptic position cue only (pVHP).

## Experiment 1

### Methods

#### Participants

Eighteen participants took part in this experiment (4 male, age 25.3 ± 8.2). All had normal or corrected-to-normal vision and no known history of neurological disorders. All of the participants were naïve to the purpose of the experiment and were provided with a subsistence allowance. The experiment was undertaken with the understanding and informed written consent of each participant and the experimental procedures were approved by the Institutional Review Board of New York University Abu Dhabi.

#### Apparatus

The set of stimuli consisted of three 3D-printed rectangular cuboids with depths of 40, 50, 60 mm, all the same height (120 mm) and width (25 mm). A chin rest was positioned at the edge of the experimental table and its height was adjusted such that the participants’ eyes were 440 mm above the table surface. During the experiment the three target objects were positioned 350 mm in the sagittal direction with respect to the table’s edge. Thus, in the peripheral vision condition, the top of the objects was at approximately 45 degrees of eccentricity with respect to the participants’ gaze (Figure 1A). This eccentricity allowed to increase the visual uncertainty without completely eliminating the availability of visual cues (Goodale & Murphy, 1997; Schlicht & Schrater, 2007). A custom-made eye-tracker was attached to the left rod of the chin rest with a locking arm (JB01291-BWW). The eye-tracker consisted of a modified webcam (Vivitar V49252), with a sampling frequency of 30 Hz and an accuracy of ±10 mm. An array of 25 infrared LEDs was positioned on the table 40 cm far and 30 cm to the left of the participant. The activation and deactivation of LEDs was controlled by an Arduino YÙN board via Matlab (Mathworks Inc, Natick, MA, USA) by a custom program, which also computed the pupil coordinates from the sampled eye images. The start position of the right hand was defined by a 5 mm high rubber bump with a diameter of 9 mm attached at the edge of the table, 450 mm to the right of the participants’ mid-line. The experiment was conducted in a dark room with the experimental table illuminated by a desk light (5W) positioned on the left side of the participant.

**Figure 1:**
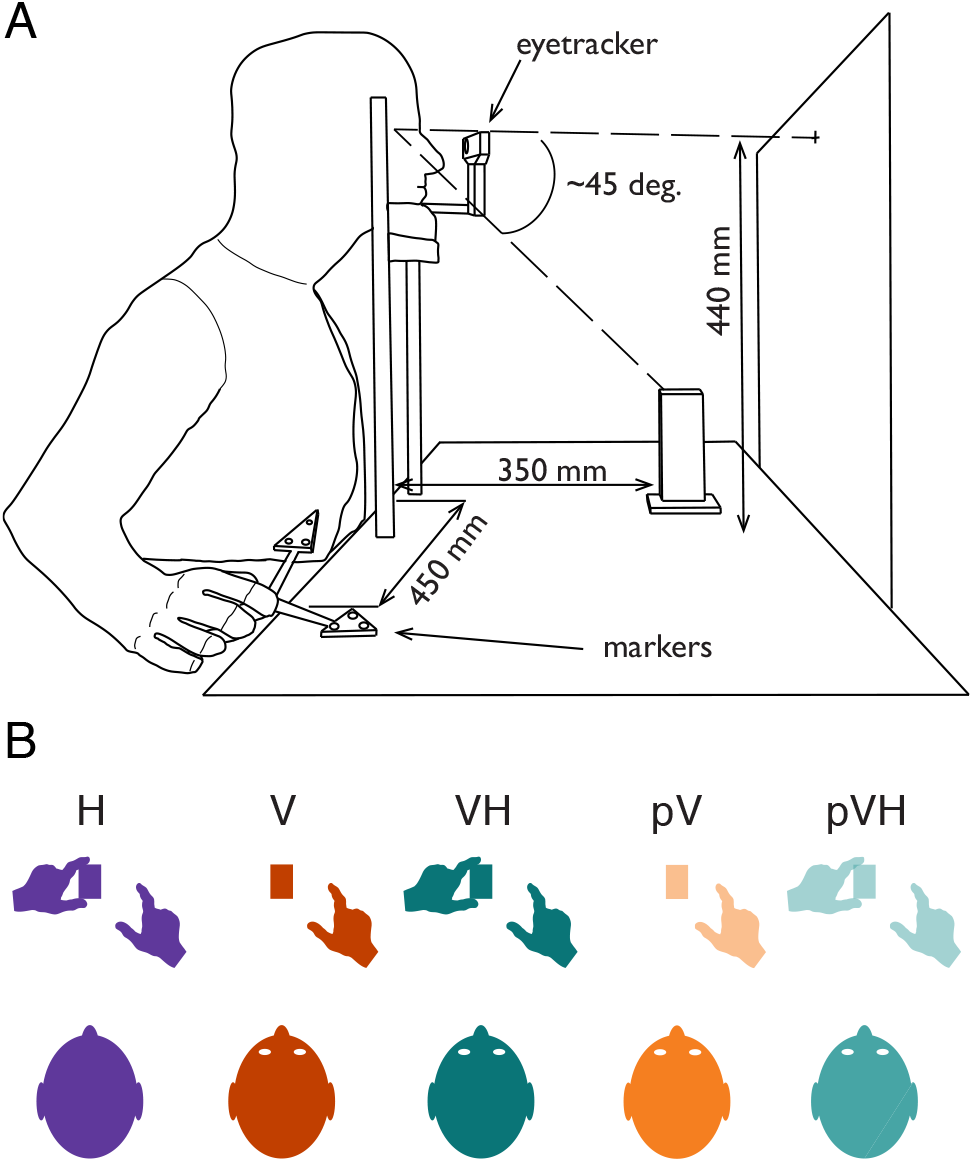
(A) Experimental setup. Participant’s head was resting on a chin rest. In the Haptic condition (H), participants were blindfolded. In the Visual (V) and Visuo-Haptic (VH) conditions, the participant’s gaze was directed toward the object. In the Visual-Peripheral (pV) and Visuo-Haptic-Peripheral (pVH) conditions, the participant was fixating a small white square on the frontoparallel board and thus the object was in peripheral vision at approximately 45 degrees of eccentricity with respect to the participant’s gaze direction. (B) Representation of the task in each condition. The grasping action was always performed with the right hand. In H and VH, the participant was already holding the object with the left hand before the start of the grasping action. In V, only vision was available. The pV and pVH conditions were identical to the V and VH conditions, but the object was seen in peripheral vision.

A black panel (600 mm wide, 500 mm high) was positioned 450 mm far from the participants’ position (i.e., behind the object). A small white square (5 mm × 5 mm) was positioned at the center of the panel, at a height of 440 mm, and acted as the fixation point in the peripheral vision block. A cardboard panel (400 mm wide, 300 mm high) was used to prevent vision of the workspace (but not of the board with the fixation point) between trials in the central and peripheral blocks, whereas a pair of occlusion goggles was used to prevent vision in the Haptic condition (Red Scientific, Salt Lake City, UT, USA). A pure tone of 1000 Hz, 100 ms length was used to signal the start of the trial, while a tone of 600 Hz with the same length was used to signal its end.

Index, thumb and wrist movements were acquired on-line at 200 Hz with sub-millimeter resolution by using an Optotrak Certus system (Northern Digital Inc., Waterloo, Ontario, Canada). The position of the tip of each digit was calculated during the system calibration phase with respect to two rigid bodies defined by three infrared-emitting diodes attached on each distal phalanx (Nicolini, Fantoni, Mancuso, Volcic, & Domini, 2014). An additional marker was attached on the styloid process of the radius to monitor the movement of the wrist. The Optotrak system was controlled by the MOTOM toolbox (Derzsi & Volcic, 2018).

#### Procedure

Participants sat comfortably at the table with their torso touching its edge. All the trials started with the thumb and index digit of the right hand positioned on the start position, the left hand positioned on the left side of the chin rest and the head on the chin rest (Figure 1A). The height of the chair was adjusted to keep the eyes at a fixed height to maintain the object at a fixed visual angle. Participants were required to perform a precision grip with their right thumb and index digit along the depth axis of the stimulus.

Before each trial, the cardboard panel was placed in front of the participant to cover the workspace, and the object was placed in its position 350 mm in front of the participant. The experimenter then removed the cardboard panel and after a variable period the start tone was delivered. The participant had to perform a right-handed reach-to-grasp action toward the object at a natural speed. No reaction time constrains were imposed. Three seconds after the start tone, the end sound was delivered, and the participant had to move the right hand back to the start position. The cardboard was then placed in front of the participant, the object was set to the new required size and the next trial started.

Five different conditions (Figure 1B) were performed: Haptic (H), Visual (V), Visuo-Haptic (VH), Peripheral Vision (pV), and Peripheral Vision plus Haptic (pVH). In the H condition, vision was prevented for the whole duration of the condition. Before each trial, the experimenter signaled to the participant to hold the object with their left hand along its depth axis (i.e., sense its size and position by means of touch and proprioception). In the V condition, as soon as the cardboard was removed the experimenter instructed the participant to look at the object which was in the central visual field (the left hand was kept on the table close to the chin rest). In the VH condition, the participant had to hold the object with their left hand and look at the object. The pV and pVH conditions were identical to the V and VH conditions except that participants were instructed to look at the fixation point instead of foveating the object, so that the target object was always in visual periphery (Figure 1A). Whereas in the pV condition only peripheral vision was available, in the pVH condition participants were asked to also hold the object with their left hand. Eye fixations in the these two conditions were monitored with the eye-tracker, which started sampling as soon as the experimenter placed the cardboard panel between the participant and the object (the cardboard height was lower than the fixation point, but high enough to cover the target object), and stopped when the end of the trial sound was delivered. If the algorithm detected an eye movement of ∼10 mm in the horizontal or vertical direction from the fixation point, the trial was discarded and repeated later in the condition. The five conditions were divided in two main experimental blocks. The H, V, and VH conditions were part of the Central vision block, whereas the pV and pVH conditions were part of the Peripheral vision block.

The Central and Peripheral vision blocks were performed in sequence, while the order of the conditions (H, V, VH, pV and pVH) was randomized within blocks and across participants. The differently sized objects were presented in a random order and ten repetitions were performed for each object size and condition, which led to a total of 150 trials per participant. In order to get accustomed with the task, participants underwent a training session of ten trials before each condition.

#### Data analysis

Kinematic data were analyzed in R (R Core Team, 2020). The raw data were smoothed and differentiated with a third-order Savitzky-Golay filter with a window size of 21 points. These filtered data were then used to compute velocities and accelerations in three-dimensional space for each digit and the wrist. Movement onset was defined as the moment of the lowest, non-repeating wrist acceleration value prior to the continuously increasing wrist acceleration values (Camponogara & Volcic, 2019b; Volcic & Domini, 2016), while the end of the grasping movement was defined on the basis of the Multiple Sources of Information method (Schot, Brenner, & Smeets, 2010). We used the criteria that the grip aperture is close to the size of the object, that the grip aperture is decreasing, that the second derivative of the grip aperture is positive, and that the velocities of the wrist, thumb and index finger are low. Moreover, the probability of a moment being the end of the movement decreased over time to capture the first instance in which the above criteria were met. Trials in which the end of the movement was not captured correctly or in which the missing marker samples could not be reconstructed using interpolation were discarded from further analysis, the exclusion of these trials (158 trials, 5.8% in total) left us with 2542 trials.

We focused our analyses on two dependent variables: the peak grip aperture, defined as the maximum Euclidean distance between the thumb and the index finger, and, the peak velocity of the hand movement, defined as the highest wrist velocity along the movement. We analyzed the data using Bayesian linear mixed-effects models, estimated using the brms package (Bürkner, 2017) which implements Bayesian multilevel models in R using the probabilistic programming language Stan (Carpenter et al., 2017). The models included as fixed-effects (predictors) the categorical variable Condition (H, V, VH, pV and pVH) in combination with the continuous variable Size. This latter was centered before being entered in the models, thus, the estimates of the Condition parameters (*β*_*Condition*_) correspond to the average performance of each Condition. The estimates of the parameter Size (*β*_*Size*_) correspond instead to the change in the dependent variables as a function of the object size. All models included independent random (group-level) effects for subjects. Models were fitted considering weakly informative prior distributions for each parameter to provide information about their plausible scale. We used Gaussian priors for the Condition fixed-effect predictor (peak grip aperture *β*_*Condition*_: mean = 90 and sd = 40; peak velocity *β*_*Condition*_: mean = 1100 and sd = 200). For the Size fixed-effect predictors we used a Cauchy prior distribution centered at 0 with a scale parameter of 2.5. For the group-level standard deviation parameters and sigmas we used Student *t*-distribution priors (peak grip aperture all sd parameters and sigma: *df* = 3, scale = 10; peak velocity all sd parameters and sigma: *df* = 3, scale = 170). Finally, we set a prior over the correlation matrix that assumes that smaller correlations are slightly more likely than larger ones (LKJ prior set to 2).

For each model we ran four Markov chains simultaneously, each for 16,000 iterations (1,000 warm-up samples to tune the MCMC sampler) with the delta parameter set to 0.9 for a total of 60,000 post-warm-up samples. Chain convergence was assessed using the 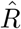 statistic (all values equal to 1) and visual inspection of the chain traces. Additionally, predictive accuracy of the fitted models was estimated with leave-one-out cross-validation by using the Pareto Smoothed Importance Sampling. All Pareto k values were below 0.5.

The posterior distributions we have obtained represent the probabilities of the parameters conditional on the priors, model and data, and, they represent our belief that the “true” parameter lies within some interval with a given probability. We summarize these posterior distributions by computing the medians and the 95% Highest Density Intervals (HDI). The 95% HDI specifies the interval that includes with a 95% probability the true value of a specific parameter. To evaluate the differences between parameters of two conditions, we have simply subtracted the posterior distributions of *β*_*Condition*_ and *β*_*Size*_ weights between specific conditions. The resulting distributions are denoted as the credible difference distributions and are again summarized by computing the medians and the 95% HDIs.

For statistical inferences about the *β*_*Size*_ we assessed the overlap of the 95% HDI with zero. A 95% HDI that does not span zero indicates that the predictor has an effect on the dependent variable. For statistical inferences about the differences of the model parameters, *β*_*Condition*_ and *β*_*Size*_, between conditions, we applied an analogous approach. A 95% HDI of the credible difference distribution that does not span zero is taken as evidence that the model parameters in the two conditions differ from each other.

## Results and Discussion

Based on our previous results (Camponogara & Volcic, 2019a, 2019b, 2021b), we expect the multisensory condition in central vision (VH) to exhibit faster grasping movements with smaller peak grip apertures than the V and H unisensory conditions. Likewise, we expect the peripheral vision conditions (pV, pVH) to show a decline in performance with respect to their corresponding central vision conditions (V, VH), because peripheral vision is characterized by a higher visual uncertainty. However, two main scenarios are considered for the peripheral vision conditions. If haptic size is largely involved in the control of grasping, we expect faster movements, with narrower peak grip apertures and a better grip aperture scaling in pVH compared to pV. If haptic size does not play a relevant role, we expect actions in pVH to be faster and with narrower peak grip apertures than in pV, but with no improvement in grip aperture scaling, that is, the sensitivity to changes in object size would be equivalent to the pV condition.

We confirmed that movements performed in central vision were faster and with a narrower peak grip aperture in multisensory compared to each unisensory conditions (Figure 2, panels A and C) (Camponogara & Volcic, 2019a, 2019b, 2021b). Interestingly, the same pattern of results was found also in peripheral vision (Figure 2, panels B and D), confirming that haptics and vision are integrated also when vision is degraded. As expected, actions were slower and were performed with a wider grip aperture in peripheral compared to central vision, in both unisensory and multisensory conditions (V vs. pV and VH vs. pVH, Figure 2). Interestingly, while the peak grip aperture scaled similarly in V and VH (Figure 2C), the scaling was stronger in pVH compared to pV (Figure 2D), suggesting a different support of haptics when acting in central and peripheral vision.

**Figure 2:**
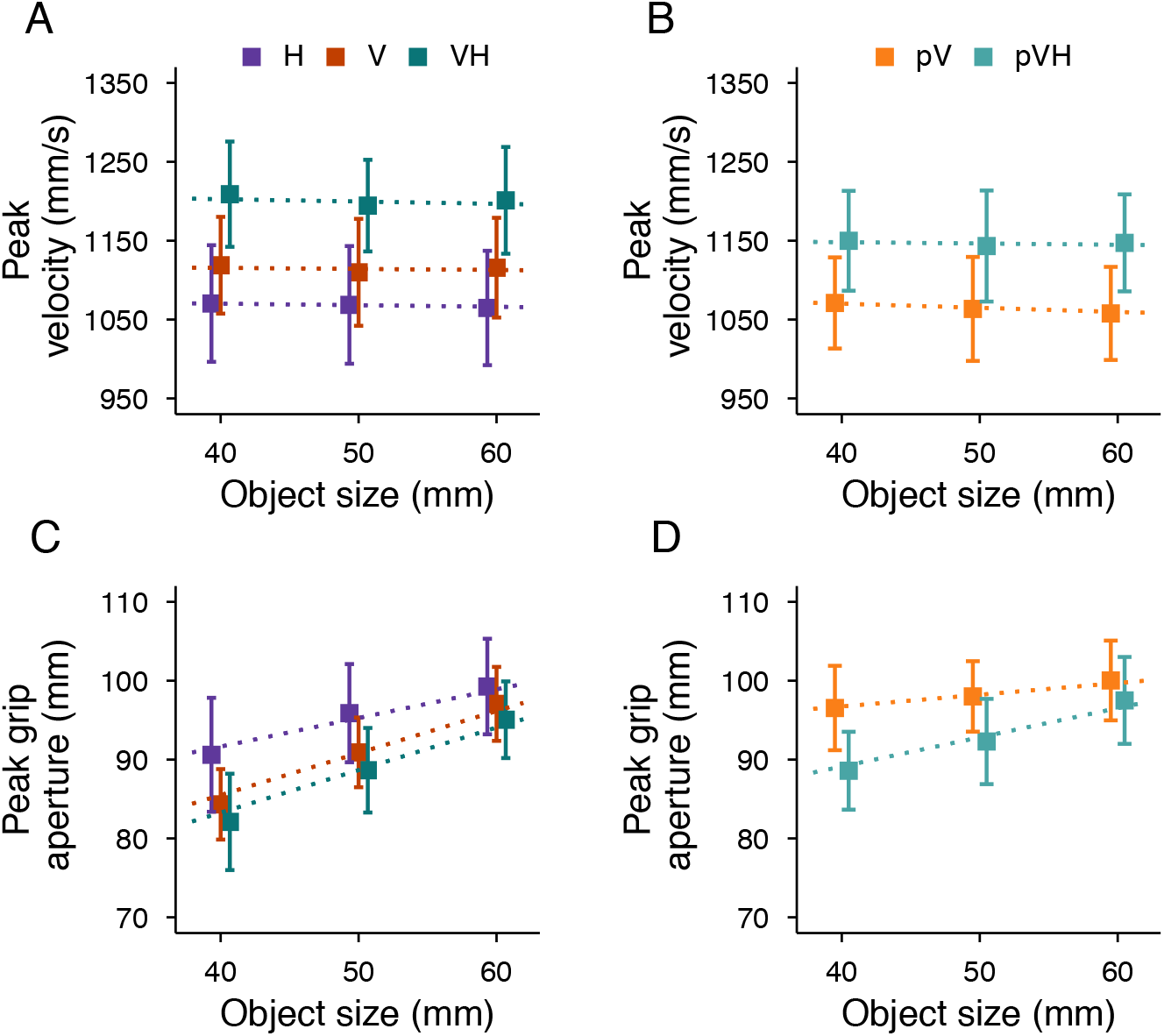
Top row: Average peak velocity as a function of the object size in the Central vision (A) and Peripheral vision (B) blocks in Experiment 1. Bottom row: Average peak grip aperture as a function of the object size in the Central vision (C) and Peripheral vision (D) blocks. Error bars represent the standard error of the mean. Dotted lines show the Bayesian mixed-effects regression model fits.

### Central vision

In central vision, the peak velocity was modulated according to the available sensory information (Figure 3A), with an advantage of multisensory over unisensory grasping, and of vision over haptics. The peak velocity was credibly higher in VH compared to V and H, and, in V compared to H (Figure 3B). The peak velocity was not affected by changes in object size in any of the conditions, with slope values ranging between −0.1 and −0.65 corresponding to minimal variations in peak velocity between the smallest and the largest object (∼10 mm/s difference equivalent to ∼1% of the average peak velocity).

**Figure 3:**
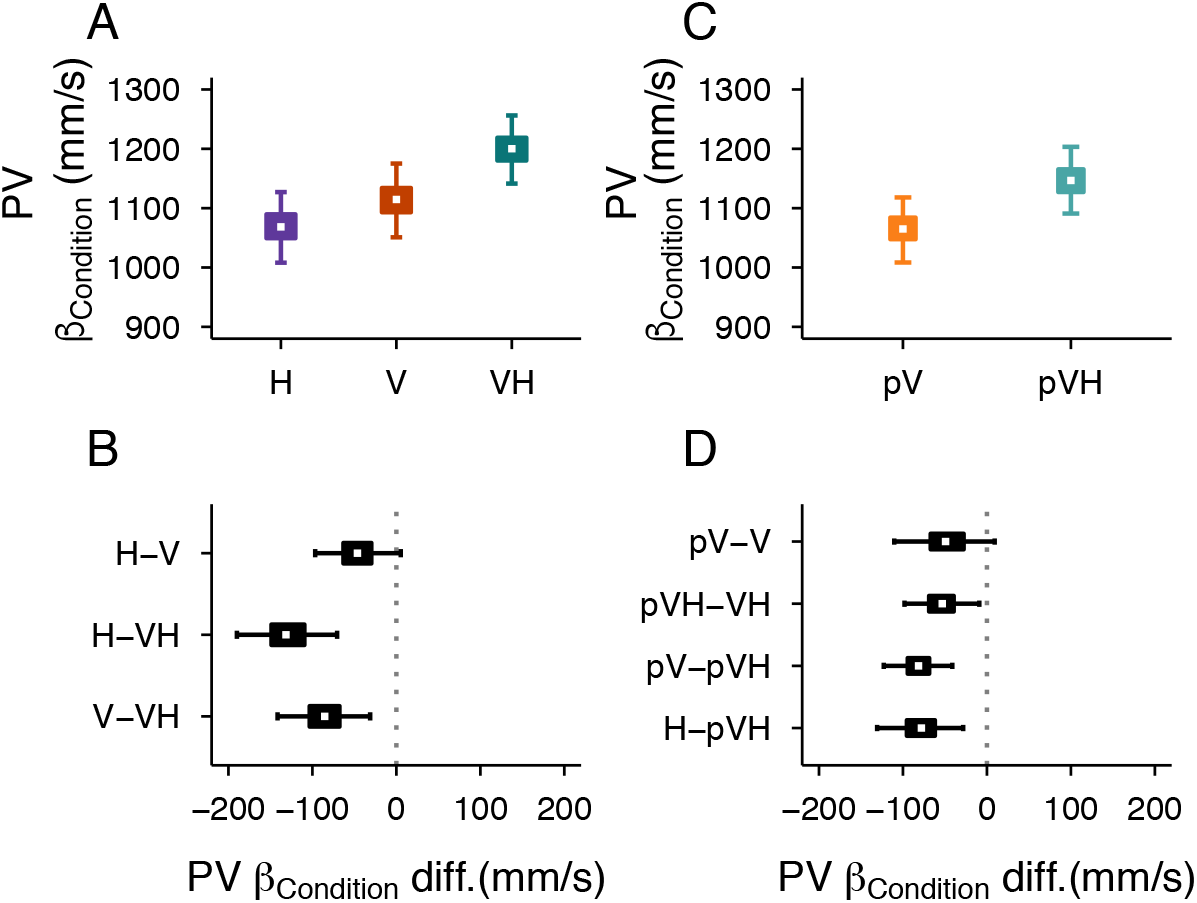
Peak velocity results (PV) in Experiment 1. Central vision block (left column): (A) Posterior beta weights of the Bayesian linear mixed-effects regression model for the predictor Condition, and (B) Credible difference distributions between conditions. Peripheral vision block (right column): (C) Posterior beta weights and (D) Credible difference distributions between conditions for the predictor Condition. White dots represent the median, the boxes represent the 50% HDIs, and the areas between whiskers represent the 95% HDIs of the distributions.

The peak grip aperture was also clearly affected by the available sensory inputs (Figure 4A). Peak grip aperture was credibly smaller in the VH condition compared to the H condition, and in V compared to the H condition (Figure 4B). A tendency of observing a credibly smaller peak grip aperture in VH compared to V condition was also observed. These results replicate our previous findings and further corroborate that the simultaneous availability of visual and haptic inputs leads to a multisensory advantage (Camponogara & Volcic, 2019b, 2021b).

**Figure 4:**
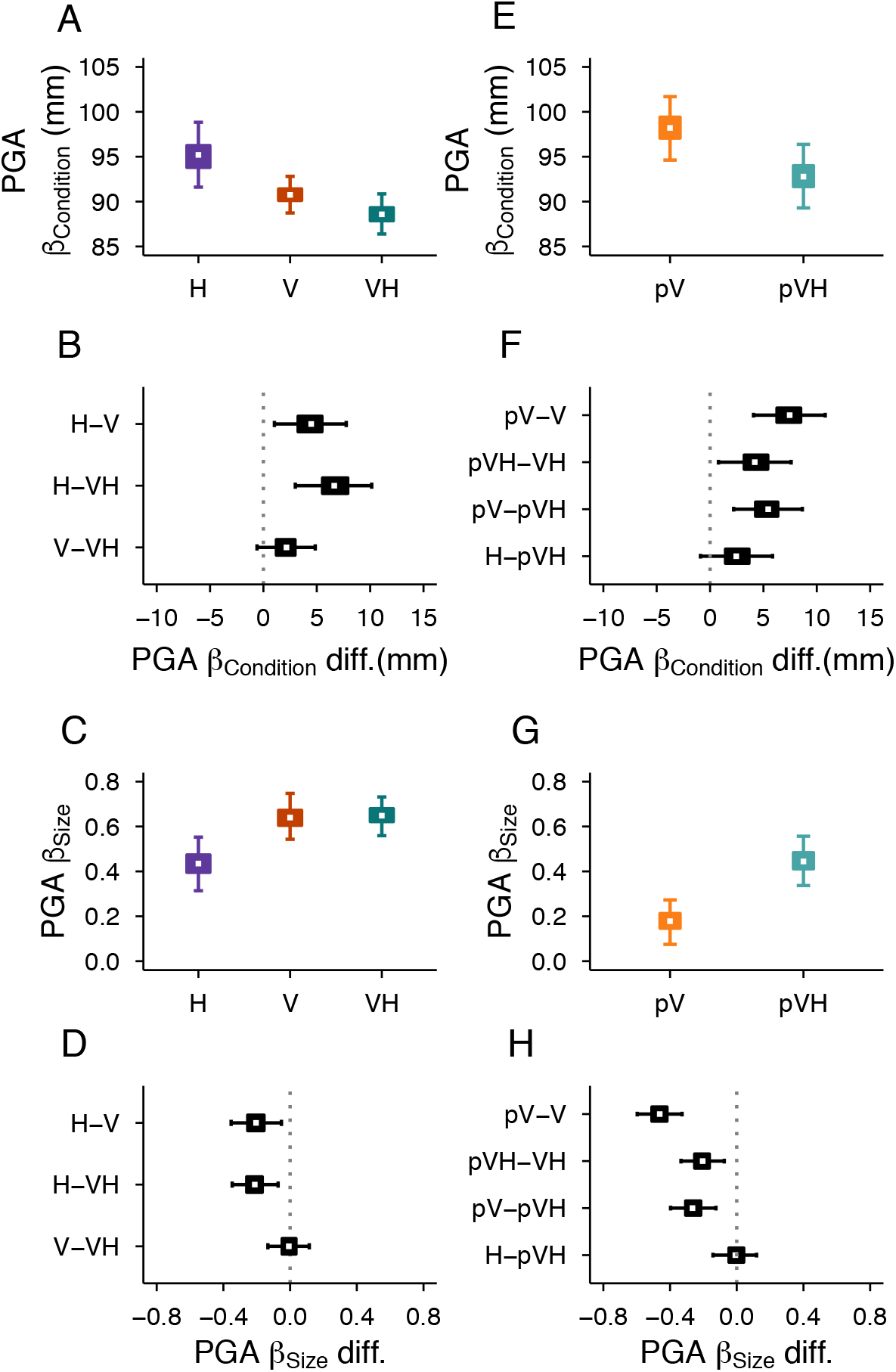
Peak grip aperture results (PGA) in Experiment 1. Central vision block (left column): Posterior beta weights of the Bayesian linear mixed-effects regression model, and the credible difference distributions between conditions for the predictors Condition (A, B) and Size (C, D). Peripheral vision block (right column): Posterior beta weights and the credible difference distributions for the predictors Condition, (E, F) and Size (G, H). White dots represent the median, the boxes represent the 50% HDIs, and the areas between whiskers represent the 95% HDIs of the distributions

The peak grip aperture scaled with object size in all conditions (Figure 4C). The scaling was equivalent in the VH and V conditions, and stronger compared to the H condition (Figure 4D). This can be considered as a sign that, in central vision, the peak grip aperture modulation in multisensory grasping is mainly based on the visual size cue, as suggested by our previous study (Camponogara & Volcic, 2021b).

### Comparisons between central and peripheral vision

Additional haptic inputs affected both peak velocity and peak grip aperture also in peripheral vision (Figure 3C and Figure 4E). As observed for central vision, holding the object with the contralateral hand facilitated faster movements and reduced grip apertures highlighting again the beneficial role of haptics.

The concurrent availability of peripheral vision and haptics enabled faster movements compared to when either of the two modalities was presented in isolation (Figure 3D, pV−pVH, H−pVH comparisons). As expected, reach-to-grasp actions toward peripherally seen objects were slower than those toward centrally seen objects. The peak velocity was credibly lower in pVH compared to VH, and there was a tendency for a credibly lower peak velocity in pV compared to V (Figure 3D, pV−V and pVH−VH comparisons).

The peak grip aperture credibly increased when the object was in peripheral compared to central vision, both with or without the support of concurrent haptic information (Figure 4F, pV−V, pVH−VH comparisons). However, the switch from central to peripheral vision increased peak grip apertures more strongly when the grasping behavior was not supported by additional haptic information (pV−V vs. pVH−VH). The effect of adding haptic information to peripheral vision resulted in credibly narrower peak grip apertures (Figure 4F, pV−pVH comparison), whereas adding peripheral vision to haptics led to only a minor improvement (Figure 4F, H−pVH comparison).

The availability of concurrent haptic size and position cues also partially prevented the typical worsening of the scaling of the grip aperture when grasping is guided only by peripheral vision (Figure 4G). Object size scaling was credibly weaker in peripheral compared to central vision (Figure 4H, pV−V, pVH−VH comparisons), but the scaling of the peak grip aperture was credibly stronger in the pVH condition compared to the pV condition (Figure 4H, pV−pVH comparison), which was, in turn, identical to the H condition (Figure 4H, H−pVH comparison). It is interesting to note that while in central vision the grip aperture scaled similarly in the unisensory visual and in the multisensory conditions (Figure 4D, V−VH), the grip aperture in visual periphery scaled more strongly in the multisensory compared to the unisensory visual condition (Figure 4H, pV−pVH). This suggests that haptic object position and size information are flexibly used according to the quality of visual information.

Figure 5A summarizes all the conditions in terms of peak velocity, peak grip aperture and the scaling of peak grip aperture as a function of object size. Conditions from worst (larger grip apertures and lower velocity) to best (smaller grip apertures and higher velocity) grasping performance lie along the diagonal line connecting the top-left to the bottom-right corners and are denoted with smaller to larger dot sizes indicating their respective slope of peak grip apertures. Two aspects are again evident here. First, both conditions with peripheral vision (pV and pVH) are inferior to their respective central vision conditions (V and VH). Second, complementing peripheral vision with haptic inputs leads to a superior grasping performance than when actions are guided only by peripheral vision (pV vs pVH). Interestingly, in peripheral vision haptics improved the grip aperture, peak velocity and the overall scaling of the peak grip aperture to a higher extent than in central vision. Figure 5 (B and C) show that these improvements in grip aperture, movement velocity and slope of grip aperture unfold along the whole movement trajectory, with a smaller grip aperture, higher velocity and a better scaling in pVH compared to pV already from the early stages of the action.

**Figure 5:**
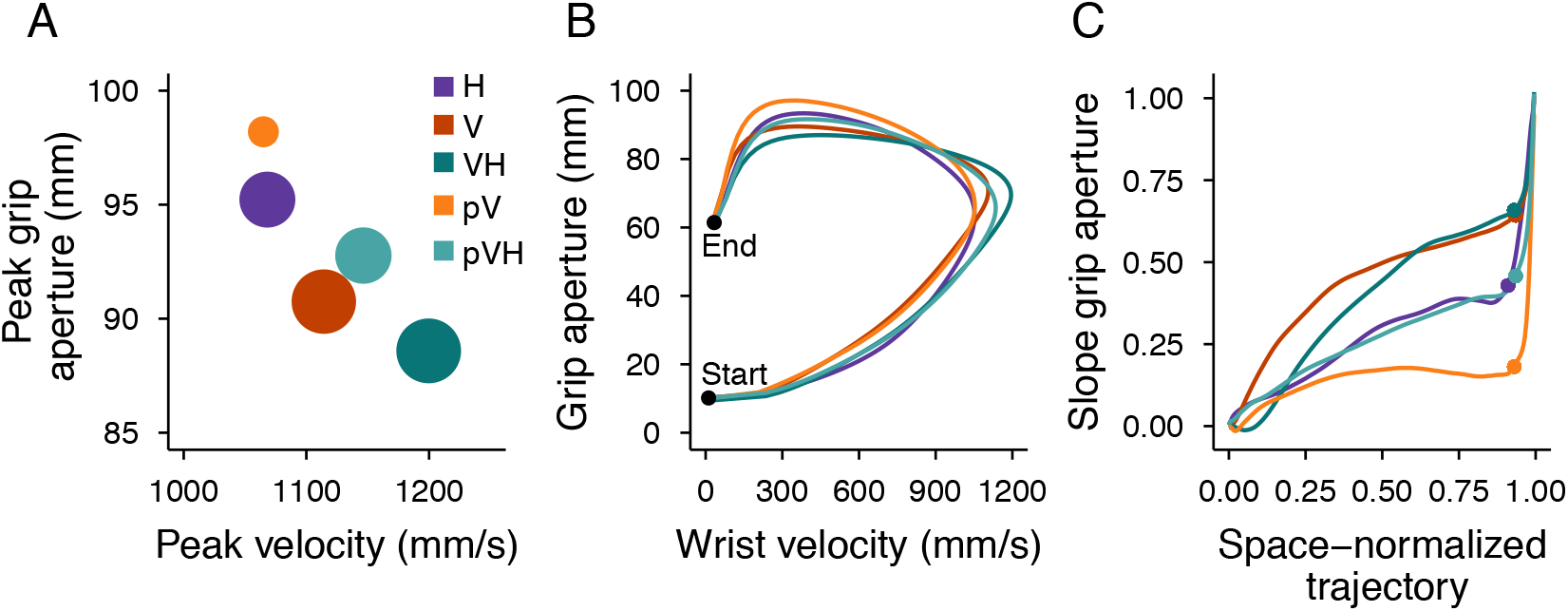
Summary of Experiment 1 results. (A) Relationship between the peak grip aperture and the peak velocity. The areas of the dots represent the slope of the peak grip aperture as a function of the object size (the higher the slope the larger the dot). (B) Relationship between the grip aperture and the wrist velocity from the start to the end of the movement. The lines representing each condition were obtained by resampling each movement trajectory in 201 steps evenly spaced along the three-dimensional path and by then averaging the grip aperture and movement velocity over all participants and sizes for each step of the space-normalized movement trajectory. (C) Slope of the grip aperture along the space-normalized trajectory. The slope values were computed by fitting a linear model with the grip aperture as a function of the object size for each step of the space-normalized trajectory. The dots represent the point of the trajectory at which the peak grip aperture occurred.

## Experiment 2

The results of Experiment 1 show that, as for central vision, actions toward handheld objects in peripheral vision are performed faster and with narrower grip apertures than those toward only (peripherally) seen objects. This suggests that visual and haptic inputs are successfully integrated even when vision is disrupted. However, the partially restored grip aperture scaling observed in peripheral multisensory grasping could have two different origins that either incorporate haptic size cues or not. If the haptic size cue is critical for hand shaping in peripheral multisensory grasping, we expect that its removal would resemble the peak grip aperture and its scaling observed in the pV condition. Instead, if the hand shaping is mainly determined by visual size cues which are improved by the availability of haptic positional information, as seen in central vision (Camponogara & Volcic, 2021b), the haptic position cue should be sufficient to attain the same level of peak grip aperture and its scaling as when all haptic cues are provided. As long as the haptic position cue is available, the presence or absence of the haptic size cue should not affect peak velocities, which should be higher than when only peripheral vision is available. To tease apart the relative contribution of these haptic inputs, we systematically manipulated the haptic size availability. In the Peripheral Vision plus Haptic Position condition (pVHP), we introduced a new set of objects which were identical to those used in Experiment 1, but had the lower half replaced by a post which did not co-vary with the size of the objects (Figure 6A). Thus, in the pVHP condition, participants were holding the post with their left hand (Figure 6B), which provided only haptic positional but no relevant size information, while simultaneously seeing the object in the periphery. This pVHP condition was performed on a new group of participants together with the Peripheral Vision (pV) and Peripheral Vision plus Haptic (pVH) conditions, which were the same as in Experiment 1.

**Figure 6:**
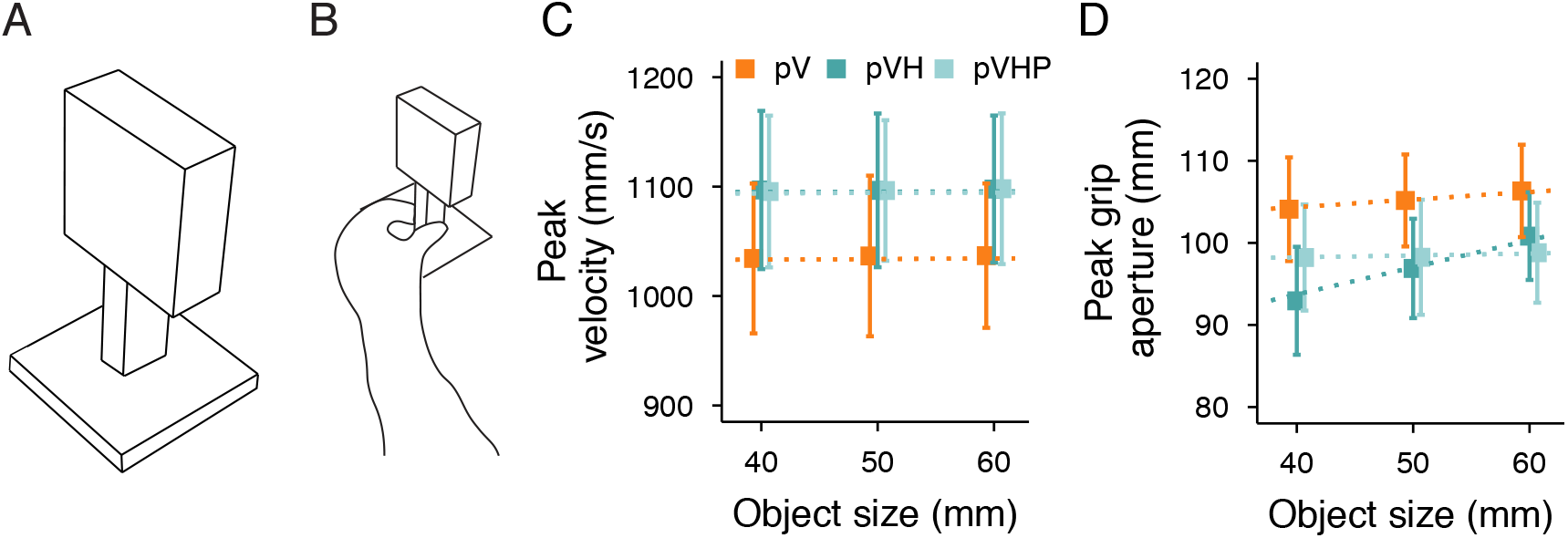
(A) Example of a stimulus used in the pVHP condition of Experiment 2. (B) Left hand holding the post during the pVHP condition. Average peak velocity (C) and peak grip aperture (D) as a function of the object size in Experiment 2. Error bars represent the standard error of the mean. Dotted lines show the Bayesian mixed-effects regression model fits.

### Methods

#### Participants

Eighteen new participants took part in Experiment 2 (6 male, age 20.7 ± 3.5). All had normal or corrected-to-normal vision and no known history of neurological disorders. All of the participants were naïve to the purpose of the experiment and were provided with a subsistence allowance. The experiment was undertaken with the understanding and informed written consent of each participant and the experimental procedures were approved by the Institutional Review Board of New York University Abu Dhabi.

#### Apparatus

The experimental setup was the same as in Experiment 1 (Figure 1A), except that two set of stimuli were used: the first set was the same as in the first experiment (Figure 1A), whereas the second set of stimuli consisted of five rectangular cuboids of 60 mm height supported by a 60 mm high post which was 10 mm deep and 25 mm wide (Figure 6A). The upper part of these stimuli was identical to the first set of stimuli and thus varied in depth across trials. The post supporting the upper part had instead a fixed depth.

#### Procedure

The procedure was the same as for the Peripheral block of Experiment 1. In the pV condition and the pVH condition the first set of objects was presented (Figure 1A). In the pVHP condition, the second sets of object was used (Figure 6A). In this case, participants held with their left hand the post that supported the target object (Figure 6B). Thus, while in the pVH condition haptic inputs were informative of both the object size and position, in the pVHP condition haptic inputs provided only positional object information. Therefore, peripheral vision was the only source of object size information.

The order of the conditions (pV, pVH, pVHP) was randomized across participants. Object sizes were randomized within each condition and 15 trials were performed for each object size and condition, which led to a total of 135 trials per participant. Before each condition, participants underwent a training session of ten trials to get accustomed with the task.

#### Data analysis

The raw data processing and the statistical analyses were identical to those of Experiment 1. Based on the same exclusion criteria, a total of 276 trials (11.3% in total) were excluded which left us with 2154 trials for the final analysis. As in Experiment 1, we focused our analyses on the peak grip aperture and the peak velocity of the hand movement. The 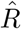 statistic and visual inspection of the chain traces confirmed successful chains convergence. All Pareto k values were below 0.5. As in Experiment 1, we report the posterior distribution of the *β*_*Condition*_ and *β*_*Size*_ for each condition, and contrast the different conditions by computing the differences between the posterior distributions for each predictor.

## Results and Discussion

Results showed that movements were performed faster and with a narrower grip aperture in the multisensory conditions (pVH, pVHP) compared to the unisensory (pV) condition. Interestingly, movements were equally fast and with a similar grip aperture either with (pVH) or without (pVHP) the haptic size cue (Figure 6C and D). However, removing the haptic size cue dramatically reduced the scaling of the grip aperture, which scaled less than when both the size and position haptic cues were available (Figure 6D).

Movements supported by haptic inputs were faster than the unisensory visual condition (Figure 7A) with a credibly higher peak velocity in pVH and in pVHP compared to pV (Figure 7B). Interestingly, as we have observed for central vision (Camponogara & Volcic, 2021b), no differences in peak velocity were found between the pVH and pVHP conditions confirming that the integration of vision and haptics is mainly concerned with the position of the object. As in Experiment 1, peak velocity was insensitive to changes in object size. The variation of the Size effect spans the [0, −0.13] range, which corresponds to a variation of the peak velocity of 2.6 mm/s from the smallest to the largest object (∼0.2% of the average peak velocity).

**Figure 7:**
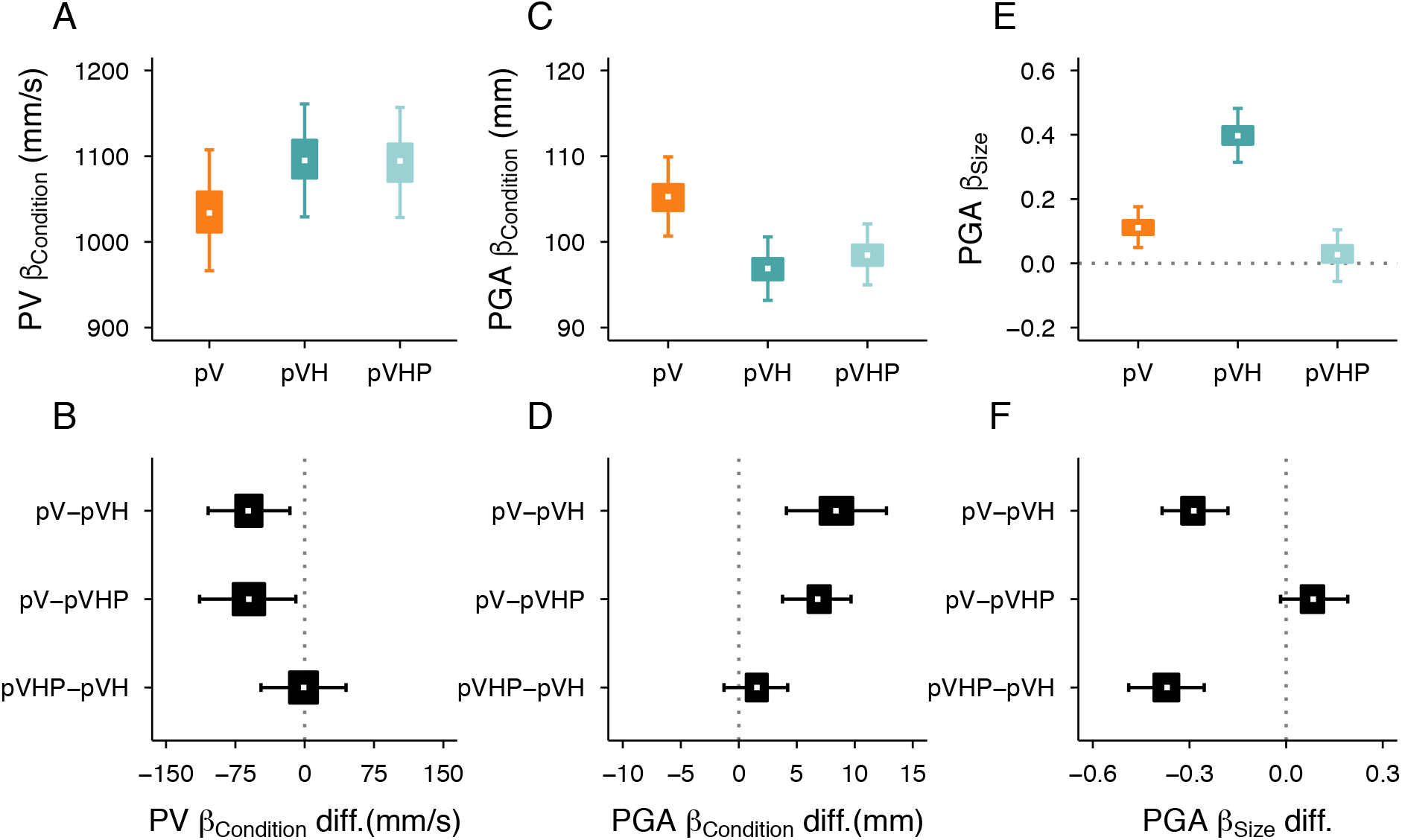
Left column: Peak velocity results (PV) in Experiment 2. (A) Posterior beta weights of the Bayesian linear mixed-effects regression model, and (B) Credible difference distributions between conditions for the predictor Condition. Middle and right columns: Peak grip aperture results (PGA) in Experiment 2. Posterior beta weights and Credible difference distributions between conditions for the predictor Condition (C, D) and Size (E, F). White dots represent the median, the boxes represent the 50% HDIs, and the areas between whiskers represent the 95% HDIs of the distributions

The analysis of the peak grip aperture reaffirmed the advantage of multisensory over unisensory conditions (Figure 7C). Peak grip aperture was credibly larger in the pV condition than in the pVH condition (Figure 7D). The peak grip aperture was also credibly larger in pV compared to pVHP, and similar between the pVH and pVHP conditions. Most importantly, providing only haptic positional information was not sufficient to accurately scale the grip aperture according to the object size (Figure 7E). We found that the peak grip aperture increased credibly less as a function of object size in pV and pVHP compared to pVH, and, it was similar between pV and pVHP conditions (Figure 7F). Thus, in degraded visual conditions the haptic positional information speeds up movements and decreases grip aperture, but haptic size is essential to modulate the grip aperture according to the object size (Figure 8A). Noticeably, the grip apertures and the movement velocities in pVH and pVHP conditions were almost indistinguishable from the beginning to the end of the movement emphasizing the specific role of the haptic position cue in improving action performance (Figure 8B). However, as can be seen in Figure 8C, the haptic size cue was crucial to refine the hand shaping aroung the object by improving the grip aperture scaling along the whole movement trajectory.

**Figure 8:**
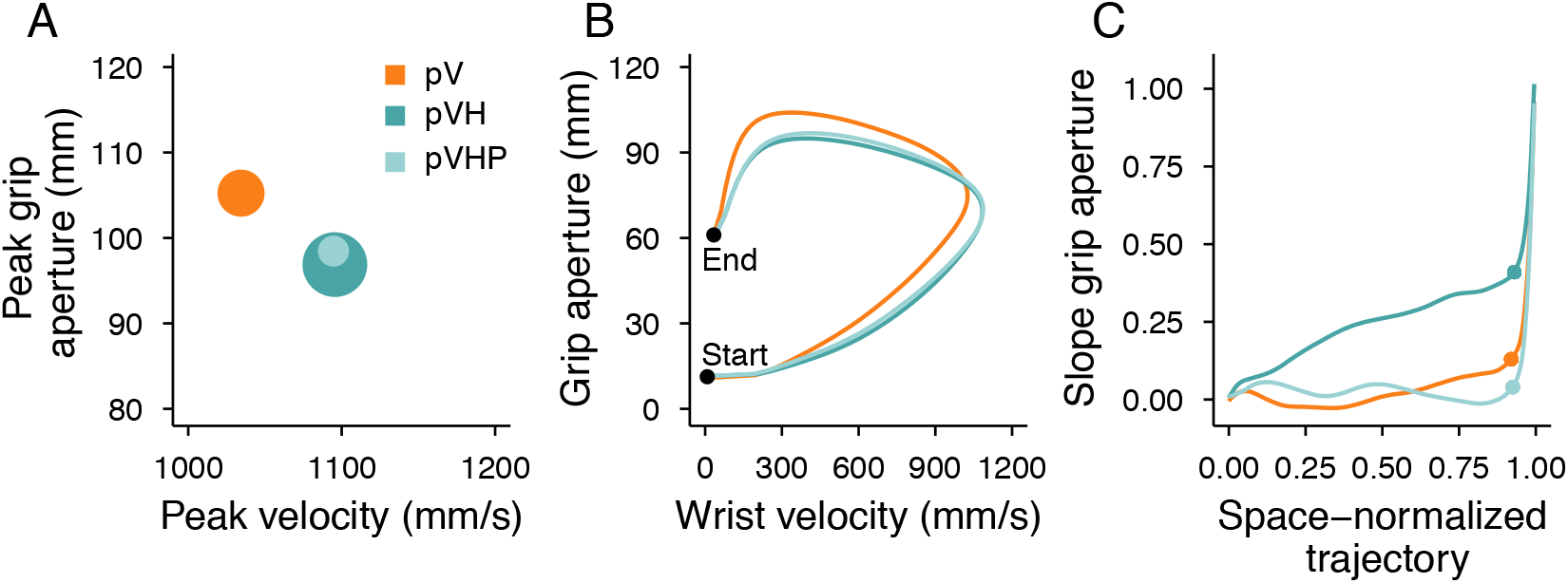
Summary of Experiment 2 results. (A) Relationship between the peak grip aperture and the peak velocity. The areas of the dots represent the slope of the peak grip aperture as a function of the object size (the higher the slope the larger the dot). (B) Relationship between the grip aperture and the wrist velocity from the start to the end of the movement. Conditions are represented by the lines obtained by resampling each movement trajectory in 201 steps evenly spaced along the three-dimensional path and by then averaging the grip aperture and movement velocity over all participants and sizes for each step of the movement trajectory. (C) Slope of the grip aperture along the space-normalized trajectory. The slope was computed by fitting a linear model with the grip aperture as dependent variable and the objects Size as factor for each step of the space-normalized trajectory. The dots represent the point of the trajectory where the peak grip aperture occurred.

## Discussion

There are two key findings of the present research. First, we found that the integration of visual and haptic object features for multisensory guided grasping occurs not only when vision is superior to haptics, but also when vision is disrupted to the extent that it becomes the less reliable modality. Second, we found that the integration of vision and haptics for multisensory guided grasping comprises both position and size cues, with the greater benefits gained by the contribution of the haptic position cue.

Visually guided grasping in central vision clearly outperformed haptically guided grasping, but it was severely degraded when vision was only peripheral. Irrespective of the quality of visual information, we have observed pronounced improvements when both vision and haptics were simultaneously available. Multisensory guided movements were faster than movements in the fastest of the unisensory conditions and grip apertures tended to be smaller than the smallest of the unisensory conditions. These findings show that the process of multisensory integration for grasping actions obeys the same rules observed in studies on visuo-haptic reaching (Camponogara & Volcic, 2021a) and visuo-haptic perception (Derrick & Dewar, 1970; Ernst & Banks, 2002; Gepshtein & Banks, 2003; Helbig & Ernst, 2007; Heller, 1983; Van Doorn et al., 2010; Wijntjes, Volcic, Pont, Koenderink, & Kappers, 2009). Thus, there is evidence in both perception and action that multisensory integration is not a rigid process in which vision simply dominates over haptics (Hay, Pick, & Ikeda, 1965; Power & Graham, 1976; Rock & Harris, 1967; Rock & Victor, 1964), but it is instead a flexible process balancing the contributions of vision and haptics depending on the quality of each source of information.

With regard to the role of the separate haptic cues, we found that enriching peripheral visual information with only the haptic position cue was sufficient to increase movement velocity and reduce grip aperture as much as when also the haptic size cue was available. It is known that the localization of objects can be strongly impaired when they are placed in visually eccentric (peripheral) positions (Bartolo et al., 2018; Bock, 1993; Henriques & Crawford, 2000; Henriques, Klier, Smith, Lowy, & Crawford, 1998). This increased positional uncertainty could be the primary cause of the worsened grasping performance usually observed when only peripheral vision is available (Brown et al., 2005; Goodale & Murphy, 1997; Hesse et al., 2012; Schlicht & Schrater, 2007; Sivak & MacKenzie, 1990, 1992; Watt et al., 2000). Our results clearly support the view that visual and haptic position cues are integrated to reduce the overall positional uncertainty, which positively influences the quality of grasping movements even when visual information is severely degraded (Camponogara & Volcic, 2021b; Chen et al., 2018). This does not exclude though that the uncertainty about object size also affects grasping movements.

The role of the haptic size cue was indeed revealed by how the grip aperture scaled according to object size. When both the haptic position and haptic size cues were provided together with peripheral vision, the scaling of the grip aperture improved with respect to the peripheral vision only condition and it was comparable to the scaling observed in the haptics only condition. This could have been an indication that the refined scaling resulted either from a reduced uncertainty about the object size driven by the availability of the haptic size cue, or, from a reduced uncertainty about the object position driven by the availability of the haptic position cue. Our results exclude the latter explanation. Providing only the haptic position cue with peripheral vision was not sufficient to induce the level of scaling observed when also the haptic size cue was available. Thus, the haptic size cue played a necessary role, because its removal, indeed, weakened the scaling of the grip aperture to the level of the peripheral vision only condition. The contributing role of the haptic size cue in reducing the overall size uncertainty is further reinforced by observing the evolution of grip aperture scaling along the whole movement trajectory. Scaling along the trajectory in multisensory peripheral vision conditions was identical to the haptic only condition when the haptic size cue was present and it was identical to the peripheral vision only condition when the haptic size cue was absent.

An additional aspect worth commenting concerns the relationship between the peak grip aperture and its scaling. The fact that peak grip aperture scales reliably with changes in object size (with a slope of approximately 0.7) is an established property of normal grasping movements (Jakobson & Goodale, 1991; Marteniuk, Leavitt, MacKenzie, & Athenes, 1990). It has also been shown that in degraded visual conditions (e.g., by removing visual feedback or by switching from binocular to monocular vision) the peak grip aperture increases and the grip aperture scaling weakens (Churchill, Hopkins, Rönnqvist, & Vogt, 2000; Hesse, Miller, & Buckingham, 2016; Keefe, Suray, & Watt, 2019; Keefe & Watt, 2009; Melmoth & Grant, 2006; Watt & Bradshaw, 2000). All our results conform with this behavior except for the multisensory condition in which only the haptic position was provided together with peripheral vision. Here the grip aperture scaling heavily decreased without the parallel increase of the peak grip aperture. This means that, if needed, the grip aperture and its scaling can be controlled independently according to the demands of a specific situation and can lead to grasping movements of generally higher quality in which collisions with objects are strategically avoided.

The associations between the visual and the haptic modality are not innate, but rather characterized by a high degree of plasticity. Vision and haptics achieve calibration during development through constant cross-sensory comparisons (Gori, Del Viva, Sandini, & Burr, 2008). Moreover, studies on cataract-treated participants showed the restoration of visual object recognition (Chen et al., 2016; Held et al., 2011) and the acquisition of multisensory integration (Senna et al., 2021) is possible within a brief period after surgery by exploiting the cross-modal interactions between vision and touch. The haptic-to-vision recalibration is also visible in adults following visuo-haptic adaptation tasks (Volcic, Fantoni, Caudek, Assad, & Domini, 2013; Wiesing, Kartashova, & Zimmermann, 2021). Our results complement these findings and raise the intriguing possibility that the haptic modality available during sensorimotor interactions with the environment could be effective in learning or restoring visuomotor functions during development and throughout the lifespan.

In sum, our results are in clear support of the view that visuo-haptic integration for grasping occurs both at the level of the position cues and at the level of the size cues confirming the hypothesis that, in optimal visual conditions, the effect of the haptic size cue is usually masked by the dominance of the more reliable visual size cue. When vision was disrupted, both haptic position and haptic size cues played a relevant role in shaping the grasping movements. It is, however, important to note that most of the advantages in multisensory grasping stem from the contribution of the haptic position cue. As we have previously suggested (Camponogara & Volcic, 2021b), a sensorimotor system can achieve greater robustness if it relies on the integration of visuo-haptic object features that systematically co-occur (e.g., position) than on features that can frequently differ between the two sensory modalities because of variations in object shape (e.g., size).

## Acknowledgments

We acknowledge the support of the NYU Abu Dhabi Research Enhancement Fund (grant RE183).

